# Robust Inference of Individualized Treatment Effect in Mendelian Randomization

**DOI:** 10.64898/2026.05.08.723855

**Authors:** Ruoxuan Wu, Xiudi Li, Feifei Xiao, Muxuan Liang

**Author notes:** Corresponding authors: Xiudi Li, Feifei Xiao, and Muxuan Liang.

## Abstract

Mendelian randomization (MR), leveraging genetic variants as instrumental variables (IVs), is widely used to draw causal conclusions in the presence of unmeasured confounding, but most MR analyses focus on average treatment effects and rely on strong assumptions. For precision medicine, the primary target is instead the individualized treatment effect (ITE); yet in MR, such effects are not point-identified under core IV assumptions, and valid inference for identifiable bounds of the ITEs is particularly challenging. In this work, we propose a robust partial identification inference framework for the identifiable bounds of the ITEs under MR allowing multiple IVs. Under minimal causal assumptions, we derive a sharp inference procedure for the bounds of ITE by adopting a multiplier bootstrap procedure with data-adaptive bootstrap distribution shifting and heterogeneous variance adjustment. In theory, we prove that the proposed method achieves nominal coverage and asymptotic sharpness. Further, we extend the procedure to tolerate possible invalid IVs under a minimal proportion rule assumption by aggregating over plausible IV subsets while preserving coverage. Simulation studies demonstrate that the proposed methods attain nominal coverage and substantially shorter intervals than existing procedures. We illustrate the framework using data from the Alzheimer’s Disease Neuroimaging Initiative to assess heterogeneous causal effects of TREM2 expression on Alzheimer’s disease risk across education-defined subgroups.

## 1 Introduction

Mendelian randomization (MR) has become a prominent causal inference approach in biomedical research. It uses genetic variants as instrumental variables (IVs) for potentially confounded exposures to study causal effects [10, 27, 31]. Most MR analyses target average treatment effects (ATEs) defined as the causal effects averaged over the entire population, but when causal effects vary across individuals, the ATE can mask important heterogeneity and be misleading for decisionmaking [2, 3, 21]. For example, in the Alzheimer’s Disease Neuroimaging Initiative (ADNI) study, the causal effect of gene expression on Alzheimer’s disease (AD) risk is plausibly modified by observed covariates such as educational attainment, indicating substantial heterogeneity in causal effects [23, 38]. These considerations motivate a focus on individualized treatment effects (ITEs) in MR, which describe how treatment effects vary across individuals rather than a single average. Many statistical learning approaches have been developed to estimate ITEs flexibly [24, 33, 39], but inference for individual-level causal effects tailored to MR remains much less developed. A valid IV requires four core IV assumptions: 1) exclusion restriction; 2) relevance; 3) unconfoundness; 4) consistency. Beyond these core IV assumptions, existing MR approaches obtaining point identified individual-level effects often require additional structural restrictions, such as effect homogeneity [6, 18, 28, 36] and no unmeasured effect modifications [1, 9, 22, 35, 37]. Such assumptions are difficult to justify in real-world applications such as the ADNI study, where unmeasured effect modifications are plausible. Without these restrictive causal assumptions, both estimation and inference for ITEs using existing methods can be biased. In this work, we focus on inferring the conditional average treatment effect (CATE), interpreted as the ITE, under minimal causal assumptions, i.e., only core IV assumptions are assumed.

To achieve this goal, we need to address several challenges. Let Δ(***x***) denote the CATE, where ***x*** is a *d*-dimensional covariate vector. First, under the core IV assumptions, Δ(***x***) is not point-identifiable but only partially identified, i.e., it can be localized to an identification region rather than a single value [5, 25, 34]. Specifically, for the *j*-th genetic variant and an individual with covariate ***x***, under core IV assumptions, we can construct identifiable functions *L*_*j*_(***x***) and *U*_*j*_(***x***) such that *L*_*j*_(***x***) ≤ Δ(***x***) ≤ *U*_*j*_(***x***). With multiple IVs, information is combined through intersection bounds of the form max_*j*_ {*L*_*j*_(***x***)} ≤ Δ(***x***) ≤ min_*j*_ {*U*_*j*_(***x***)}. Under partial identification, it is impossible to identify the exact CATE, but we can still learn about the direction of causal effects. For example, if max_*j*_ {*L*_*j*_(***x***)} *>* 0, then we know that CATE must be positive. Thus, it is important to construct a powerful inference procedure such that positive or negative causal effects can be detected given limited sample sizes. Second, existing methods for the inference for these intersection bounds are not tailored to MR studies. The precision-corrected procedure of [8] provides a general framework for intersection bound inference, especially when the estimators for *L*_*j*_(***x***)’s or *U*_*j*_(***x***)’s have heterogeneous variances. However, in MR analysis, the resulting intervals can be quite conservative and substantially wider than needed in finite samples. In parallel, [13] study how one can asymptotically select maximum/minimum and obtain valid and asymptotically sharp inference, but they do not account for heterogeneous variances that are common for multiple genetic variants. Third, when the IV assumptions are not satisfied for all candidate IVs, i.e., there might be invalid genetic variants, it is also unclear how to develop an inference framework robust to the existence of such invalid IVs. Overall, none of the existing methods are tailored to the inference of intersection bounds under MR: an ideal approach should simultaneously achieve asymptotically sharp inference, address heterogeneous variances, and remain robust when invalid IVs exist.

In this work, we propose robust inference procedures for ITEs using MR under minimal assumptions. First, we propose an efficient inference method based on nonparametric kernel regression for the intersection bounds. To infer these intersection bounds, e.g., max_*j*_ {*L*_*j*_(***x***)}, the kernel regression with undersmoothing can provide an asymptotic normal estimator for each *L*_*j*_(***x***); however, the key challenge is that we do not know which term will achieve the maximum/minimum. Thus, the inference procedure needs to account for this uncertainty in selecting the maximizers/minimizers while retaining efficiency. The proposed method asymptotically selects the maximizers/minimizers through adaptively shifting bootstrap distribution based on the kernel estimates to achieve valid confidence intervals, and adaptively adjusts each kernel estimates based on their variances for efficiency. In theory, we show that the resulting confidence intervals achieve the nominal coverage for the intersection bounds and are asymptotically sharp. When compared with existing methods for intersection bound inference, our proposed method often yields minimal difference against the true intersection bounds, especially when the variances are highly heterogeneous. Second, when there are potentially invalid IVs, by assuming a lower bound on the percentage of valid IVs, we propose an enumeration approach based on the proposed intersection bound inference method, which remains valid under possible invalid IVs.

This paper is organized as follows. In Section 2, we introduce the problem setup and our proposed inference framework under both valid and invalid IVs. In Section 3, we provide theoretical results establishing the validity of our methods. In Section 4, we demonstrate the superior performance of our proposed method compared with other existing methods through simulation studies under finite samples. In Section 5, we apply our method in the ADNI dataset to study the effect of gene expression on AD risk. Section 6 provides a discussion.

## 2 Methods

In this section, we propose an inference method to construct a valid interval estimation for ITE. Let ***X*** ∈ *χ* ⊆ ℝ^*d*^ be a *d*-dimensional covariate vector, *T* be a binary exposure (e.g., the expression level of a gene of interest - low vs. high), and *Y* be the observed outcome of interest. For the ease of presentation, we assume that the covariates are continuous. In the main text, we also assume binary outcomes; extension to continuous outcomes can be found in the online Supplementary Information. Under the Neyman-Rubin framework [17, 30], we introduce two potential outcomes corresponding to the outcomes if a patient receives *T* = 1 and *T* = 0, denoted as *Y*_1_ and *Y*_0_, respectively. In this work, we are interested in inferring the ITE defined as Δ(***x***) ≡ *E*[*Y*_1_ − *Y*_0_ | ***X*** = ***x***]. We will first introduce the partial identification under MR with all valid IVs and discuss the challenges in inferring ITE in this framework; then we will introduce the proposed inference method and its extension assuming only an upper bound on the percentage of potentially invalid IVs.

### 2.1 Mendelian Randomization and Partial Identification

MR uses genetic variants as IVs. Let 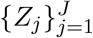 be *J* candidate genetic variants. While in many applications *Z*_*j*_’s are binary, here we also allow *Z*_*j*_’s with more than two categories, e.g., we may combine two binary generic variants and construct an IV with four categories. Among these candidate genetic variants, invalid IVs may be included. Under the Neyman-Rubin framework, we define *T*_***z***_ as the potential gene expression level given genetic variants’ profile ***Z*** = ***z***, and *Y*_***z***,*a*_ as the potential outcome given the genetic variants profile ***Z*** = ***z*** and exposure *T* = *a*. When all IVs are valid, we assume the following core IV assumptions:

1. Exclusion restriction: 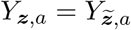 for any ***z*** and 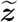, and thus we can use *Y*_*a*_ to denote them;
2. Relevance: 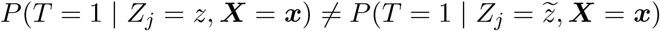 for all ***x***;
3. Unconfoundedness: (*Y*_*a*_, *T*_*z*_) ⊥ (*Z*_1_, …, *Z*_*J*_) | ***X***;
4. Consistency: ***Z*** = ***z*** implies that *T* = *T*_***z***_; further, *T* = *a* implies that *Y* = *Y*_*a*_

Under only these core IV assumptions, the ITE is not point-identifiable. Instead, from each valid IV, we can identify lower and upper bounds of ITE. That is, given a patient with covariate information ***x***, we can identify *L*_*j*_(***x***) and *U*_*j*_(***x***) such that *L*_*j*_(***x***) ≤ Δ(***x***) ≤ *U*_*j*_(***x***). This result is referred to as the partial identification of ITE in existing literature (see [4, 5, 25, 26, 34]). Here, we adopt the Balke-Pearl bound (BP bound) for binary IVs, exposure, and outcome. In BP bound, *L*_*j*_(***x***) and *U*_*j*_(***x***) are the maximum and minimum of linear combinations of conditional probabilities taking the form *P* (*Y* = *y, T* = *l* | *Z*_*j*_ = *z, X* = ***x***) for some value (*y, l, z*) ∈ {0, 1}^3^. Let *l*_*k*_ and *u*_*k*_ denote the *k*-th linear functionals in the maximum and minimum comprising the BP bound, respectively (see detailed formula in the online Supplementary Information). Note that there are *K* linear functionals in total in both the maximum and minimum, none of which depend on *j*. Let ***P***_*j*_(***x***) denote a vector composed of the conditional probabilities *P* (*Y* = *y, T* = *l* | *Z*_*j*_ = *z, X* = ***x***) when we vary the values *y, l*, and *z*. When the *j*-th IV is valid, following the BP bound, we can identify a bound for Δ(***x***), max_*k∈*[*K*]_ {*l*_*k*_ ∘ ***P***_*j*_(***x***)} ≤ Δ(***x***) ≤ min_*k∈*[*K*]_ {*u*_*k*_ ∘ ***P***_*j*_(***x***)}. Using one valid IV, the resulting bound for Δ(***x***) may be loose; by using multiple valid IVs, we can improve this bound by intersection, i.e.,

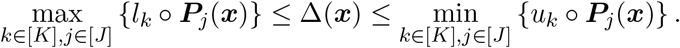

In this work, our goal is to infer *L*(***x***) and *U* (***x***) defined as

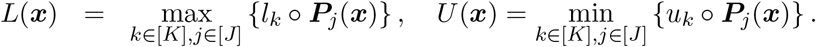

For simplicity of the notations, given an ***x*** value of interest, we will sometimes omit ***x*** in *L*(***x***) and *U* (***x***), and refer to them as *L* and *U*. We aim to construct a valid lower confidence bound (LCB) for *L* and an upper confidence bound (UCB) for *U*, denoted by 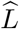 and 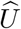 respectively, such that

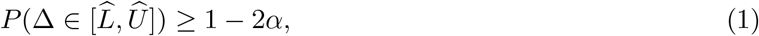

where 1 − 2*α* is the pre-specified nominal coverage. The key challenge is how to construct these *L* and *U* such that they can be as close to 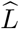 and *Û* as possible under (1).

These interval estimators provide a foundation for statistical testing under partial identification and personalized decision-making. Specifically, the constructed 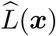 and *Û* (***x***) enable hypothesis testing about the sign, magnitude, or heterogeneity of ITEs. For example, a one-sided test of *H*_0_ : Δ(***x***) ≤ 0 versus *H*_1_ : Δ(***x***) *>* 0 rejects the null when 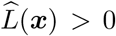, while a two-sided test of *H*_0_ : Δ(***x***) = 0 rejects when 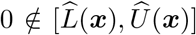. Thus, the inference of *L*(***x***) and *U* (***x***) can inform whether Δ(***x***) is positive or negative, and thus guide treatment decisions and scientific discovery.

#### Remark 1.

*When multiple valid IVs are available, the intersection bounds L*(***x***) *and U* (***x***) *allow a potentially increasing number of binary IVs. In principle, instead of intersecting bounds, our framework can also be applied when the IVs are stacked and treated as a single categorical vector* 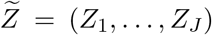. *However, this alternative requires estimating conditional probabilities of the form* 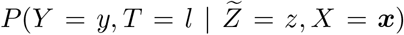 *for each joint IV configuration z. As the number of IVs J increases, the number of such configurations grows rapidly, making nonparametric estimation of these conditional probabilities statistically and computationally challenging. By working with intersection bounds across individual binary IVs, we can accommodate a large, potentially increasing number of IVs*.

### 2.2 Proposed Intersection Bound Inference

In this section, we introduce our proposed inference procedure to construct confidence bounds for *L* and *U* assuming all IVs are valid. We take *L* as an example; the inference procedure for *U* can be derived similarly.

To avoid possible model misspecification, our proposed intersection bound inference is based on nonparametric kernel estimators. Specifically, let 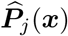 be the vector composed of

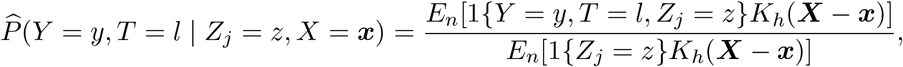

where *E*_*n*_[·] is the empirical average over *n* training samples, *K*_*h*_(·) = *h*^*−d*^*K*(·*/h*), *K* : ℝ^*d*^ → ℝ is a kernel function, and *h* is the kernel bandwidth. An initial bandwidth can be chosen via cross-validation, and we choose the final bandwidth as this initial bandwidth multiplied by an additional factor of (log *n*)^*−*1^ to ensure under-smoothing. Using an appropriate bandwidth, based on this non-parametric kernel estimator, we can construct estimator for *L*(***x***) by 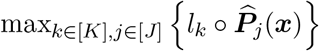. Each component 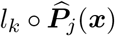 is asymptotically normal.

Based on the constructed estimators, a naive procedure would be to first select 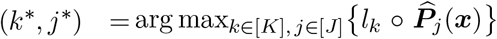, and then construct a confidence interval based on 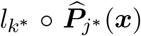. However, this procedure ignores the sampling variability in the selection step: (*k*^*\**^, *j*^*\**^) is itself data-dependent, and the maximizer of 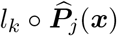 need not coincide with the maximizer of *l*_*k*_ ∘ ***P***_*j*_(***x***). Treating (*k*^*\**^, *j*^*\**^) as if it were fixed typically leads to undercoverage, as is often seen in post-selection inference problems.

To account for the randomness in (*k*^*\**^, *j*^*\**^), motivated by [13], we consider the following resampling-based procedure. Firstly, we use multiplier bootstrap to characterize the sampling randomness of 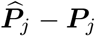 for fixed *j*. Specifically, let *W* be a random variable following a standard normal distribution independent of (*Y, T, Z*, ***X***). We construct the bootstrap random variable 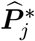, which is a vector composed of

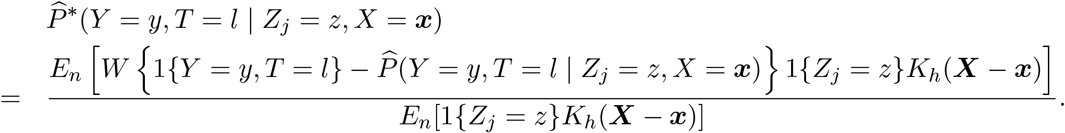

The distribution of the bootstrapped random variable 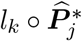 conditional on the observed sample of (*Y, T, Z*, ***X***) encodes the sampling randomness of 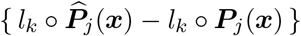. Then, we calculate the (1 − *α*)-th quantile of

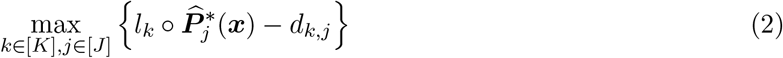

conditional on the observed sample, where

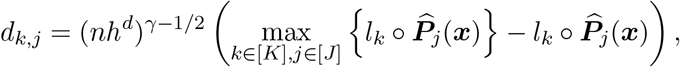

for some *γ* ∈ (0, 1*/*2). In (2), *d* is employed to shift the bootstrap distribution of 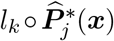 based on how far it is compared with the estimated maximum, i.e., 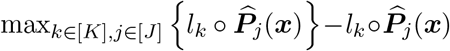. Given (*k, j*), if the value of 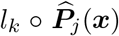 is much smaller than the maximum, then a larger shift is applied on 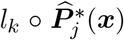. When *nh*^*d*^ +, if (*k, j*) indeed corresponds to the maxima, then 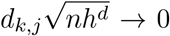; otherwise, 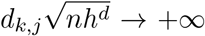. In practice, we can approximate the quantile in (2) up to arbitrary precision by computing the corresponding empirical quantile over a large number of realized multipliers ***w***.

Finally, we denote the calculated quantile as 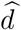, and our final LCB 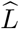 is then defined as

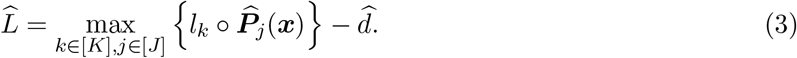

We expect that this LCB provides a valid inference for *L*, i.e., 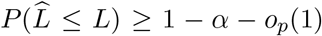; thus, a valid confidence interval for *L* can be constructed as 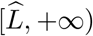. Further, the proposed inference procedure is asymptotically sharp, i.e., 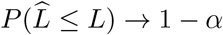 when *nh*^*d*^ → +∞.

Although the LCB 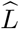 is asymptotically sharp with asymptotic size 1 − *α*, it does not mean that 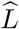 is the most efficient one among all alternatives that achieve the coverage requirement in (1). When 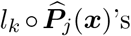 have heterogeneous variances, it is possible to construct an LCB closer to *L* that remains asymptotically sharp. The intuition is that in (3), the adjustment 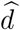 is derived based on (2) and applied to each 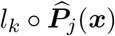 without considering possible heterogeneous variances of 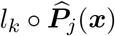. Suppose that the asymptotic variance of 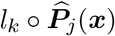 for some (*k, j*) is much larger than the others, then the value of 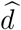 may be dominated by such (*k, j*) and unnecessarily large compared with a case where the asymptotic variances are equal. This motivates us to consider using differential adjustments based on the asymptotic variances of each term.

Motivated by [8], we consider the following procedure that accounts for possibly heterogeneous variances, which leads to an LCB that imposes the desired differential adjustments. Firstly, we construct the bootstrapped variable 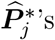 and further estimate the sample variances of 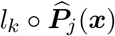 for each *k* and *j*, and denote them as 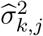. Then, instead of (2), we consider the standardized

sampling errors, i.e.,

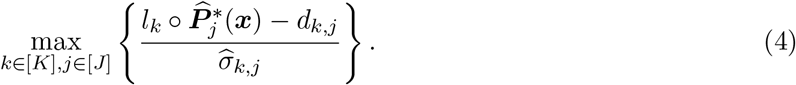

Finally, we select the (1 − *α*)-th quantile of the standardized sampling errors and denote it as 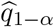. Our final LCB 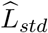 is then defined as 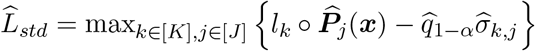. Compared with 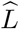 which imposes a common adjustment 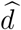 on each term without considering possible heterogeneous variances, the proposed LCB 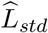 applies differential adjustments proportional to the asymptotic standard deviation of each term, and thus may lead to a shorter interval length, i.e., 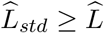.

#### Remark 2.

*Likewise, we can construct Û and Û* _*std*_ *such that P* (*Û* ≥ *U*) → 1 − *α and P* (*Û* _*std*_ ≥ *U*) → 1 − *α when nh*^*d*^ → +∞. *Combining with the inference for L, we can provide a confidence interval for* Δ(***x***) *such that*

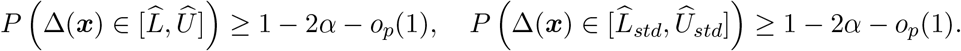

*Besides providing confidence intervals for* Δ(***x***), *the result can also be used to test whether there exists invalid IVs among* ***Z***. *Specifically, if* 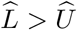 *or* 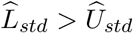, *we can reject the null hypothesis H*_0_ *that all IVs are valid with a Type I error controlled asymptotically at* 2*α*, which is equivalent to testing the validity of IVs through IV inequalities under binary IVs, treatment, and outcome [20, 29].

### 2.3 Robust Inference under Possible Invalid IVs

In this section, based on our proposed inference procedure, we propose a robust inference framework when possible invalid IVs exist. Our approach aggregates inference over candidate IV subsets and remains valid without requiring knowledge of which IVs are valid. In particular, we adopt a minimal proportion rule assumption, which posits that more than *p*% of the candidate genetic variants are valid IVs, thereby permitting a proportion of invalid IVs. This assumption assumes that the proportion of invalid IVs cannot exceed 1 − *p*%, where *p* is a pre-specified parameter. In practice, the validity of IVs may not be directly verified through the observed data, and thus the specification of *p* depends on domain knowledge. Another point of view on how to specify *p* is to treat it as a sensitivity analysis parameter to examine how inference results change with respect to the different specifications of *p*. As a special case, taking *p* = 50% recovers the majority-valid IV assumption that is commonly used in the literature [14, 15, 19].

The key challenge is that the identities of the valid IVs are unknown. Although we know a minimal proportion of valid IVs, given a single IV, in general, we do not know whether it is valid or not. Thus, to exploit the minimal proportion rule assumption, we enumerate all possible subsets of IVs containing at least *p*% of all candidate IVs; then, for each subset, we treat all IVs contained in this subset as valid (though they may not be) and construct an inference procedure for *L* using the proposed inference procedure. Under the minimal proportion rule assumption, there exists at least one subset which includes only valid IVs. Motivated by this, we take the union of the constructed intervals over all possible subsets of IVs as the final interval/inference for *L*. This procedure is guaranteed to achieve the nominal coverage for *L* as long as the minimal proportion rule assumption holds.

More formally, suppose that among *J* candidate genetic variants, a proportion *p*% are valid, which implies there exist at least *p*% × *J* valid IVs. Because the identities of the invalid IVs are unknown, we proceed conservatively by considering all possible combinations of valid IVs and this leads to 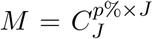 subsets. Let *ℳ* be the collection of all such subsets of IVs. For each subset *m* ∈ *ℳ*, we treat the IVs incl {uded in the subset as v}alid and conduct the proposed inference procedure, i.e., 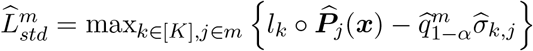, where 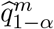 is the critical value under the subset *m*. By the minimal proportion rule assumption, at least one subset *m* must contain only valid IVs, and hence its bound is guaranteed to cover the truth with high probability. To obtain a valid overall estimate, we take the union across all candidate subsets: 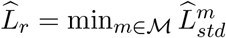. This construction ensures that the final interval 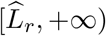 retains coverage as long as the minimal proportion rule assumption holds.

## 3 Theory

In this section, we provide theoretical guarantees for the proposed inference methods. We focus on the proposed method with all IVs valid and show that 1) the proposed interval estimator achieves the nominal coverage and is asymptotically sharp and 2) the proposed method often yields shorter interval compared with existing methods. In the online Supplementary Information, we provide a theoretical justification that motivates us to propose a tuning procedure to determine *γ*. For our proposed method, we assume the following assumptions.

### Assumption 1.

*The kernel function K*(·) *is bounded and has bounded first and second order derivatives. Denote the order of the kernel function as v; we require that v* ≥ 2.

### Assumption 2.

*The covariate vector* ***X*** ∈ *χ*, *where χ is a bounded subset of* ℝ^*d*^, *and the density function of* ***X*** *is at least v-th order continuously differentiable. In addition*, ***P***_*j*_(***x***) *is also at least v-th order continuously differentiable, and the probability and its (high-order) gradient can be uniformly bounded. Furthermore, we assume that σ*_*k,j*_ *are uniformly bounded away from* 0 *and* +∞. *We require that* 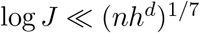 and 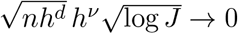.

Assumptions 1 and 2 are common assumptions for the asymptotic normality of kernel regression estimators. Under Assumptions 1 and 2, we choose *h*≪ *n*^*−*1*/*(2*v*+*d*)^ (for a fixed *J*); with such a choice of *h*, our kernel estimators satisfy that 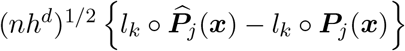 is asymptotically normal and has a sub-Gaussian tail [11, 16]. For a growing *J*, we require an even larger bandwidth to ensure the validity of high-dimensional central limit theorem as in [7]. In addition, we require the following assumption.

### Assumption 3.

*Let H* = {(*k, j*) : *l*_*k*_ ∘ ***P***_*j*_(***x***) = *L*} *and* 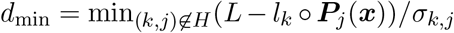. *We require that* 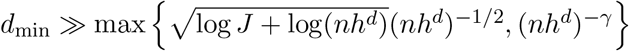.

Assumption 3 assumes that those (*k, j*)’s in *H* can be separated from other terms when *n* → +∞ under appropriate choice of *γ*. When *J* is finite and not increasing with *n*, this assumption is satisfied with any *γ* ∈ (0, 1*/*2) since *d*_min_ is finite and does not depend on *n*. However, when *J* depends on *n* and grows to infinity with *n*, it is possible that *d*_min_ decreases to 0 as *J* → +∞. In this case, Assumption 3 requires that *d*_min_ cannot approach 0 too fast in order to achieve asymptotic sharpness and tail control, especially when *J* → +∞.

Under Assumptions 1 - 3, we have the following theorem, which shows that the proposed inference method is asymptotically sharp.

### Theorem 1.

*Under Assumptions 1 - 3 and assuming all IVs are valid, for our proposed inference method in Section 2.2, we have* 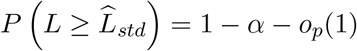.

From Theorem 1, we can also obtain the validity of the inference procedure in Section 2.3 under the minimal proportion assumption. Specifically, let *ℳ*^*\**^ ⊂ *ℳ* be the sets containing only valid IVs. Suppose that the assumptions in Theorem 1 hold for any *m*^*\**^ ∈ *ℳ*^*\**^, then the result in Theorem 1 implies that 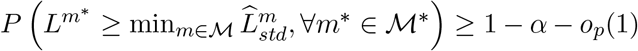; thus, as *m*^*\**^ contains only valid IVs, we have that 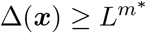.

Next, we provide a comparison with other existing methods in terms of the interval lengths. We compare our proposed LCB 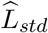 with other two competitors, namely 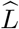, which is constructed without considering heterogeneous variances, and 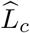 obtained via the same procedure as our proposed method but with *d*_*k,j*_ set to 0 for all (*k, j*)’s. In particular, 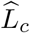 corresponds to the method in [8] without the screening procedure. Since the confidence intervals of *L* is one-sided, to compare different methods, we only need to compare 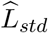 with 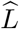 or 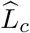.

### Theorem 2.

*For any fixed α, we have the following results: 1) We have that* 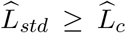 *with probability* 1; *2) Define* 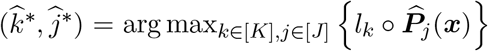. *If* 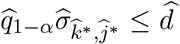 *then we have that* 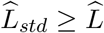.

Theorem 2 shows that compared with 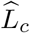, the introduction of *d*_*k,j*_ can always shorten interval lengths; in addition, we provide a sufficient condition under which 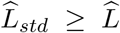. The condition 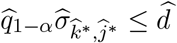 is often satisfied when there are heterogeneous variances, especially when the 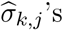 for (*k, j*) ∉ *H* are relatively large compared with those for (*k, j*) ∈ *H*. When this happens, 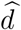 can be dominated by 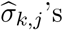 for (*k, j*) ∉ *H*, which is expected to be larger than 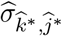 (if 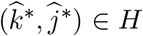); and thus, we have that 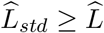.

## 4 Simulations

In this section, we conduct simulations to evaluate the finite-sample performance of our methods. We consider both the case where all IVs are valid and the case where invalid IVs exist. Through these simulations, we demonstrate that our method is more efficient compared to other methods and robust in the presence of invalid IVs.

To generate data, we first generate an unmeasured confounder *U* following a uniform distribution on [−1, 1]. Then we generate IVs depending on whether the IV is valid or not. If an IV is valid, we generate the IV following a Bernoulli distribution with a probability of 0.5 and independent from *U*; if an IV is invalid, we then generate the IV from a Bernoulli distribution with a probability of *expit* {*U/*2}. After generating the IVs independently, we generate two covariates following uniform distributions on [−1, 1]. Based on the generated IVs and covariates, we further generate the treatment *T* by a Bernoulli distribution with a probability of *T* = 1 following 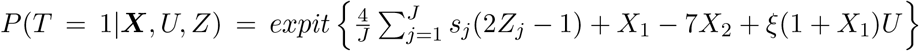, where *s*_*j*_ controls the strengths of *Z*’s, and *ξ* controls the strength of confounding involved in *T*. The outcome Y is defined as *Y* = **1**(*T* = 1) *Y*_1_ + **1**(*T* = 0) *Y*_0_, where *Y*_1_ and *Y*_0_ are potential outcomes that are generated independently using logistic regression models following *P* (*Y*_*a*_ = 1|***X***, *U*) = *expit* {*g*_1_(***X***, *U*) + (2*a* − 1)*g*_2_(***X***, *U*)} for *a* ∈ {0, 1}. In our simulations, we consider two options for *g*_1_(***X***, *U*) and *g*_2_(***X***, *U*):

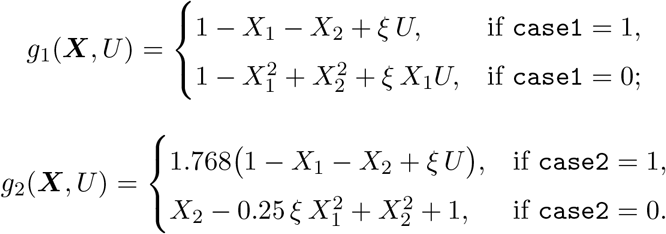

In these logistic regression models, *g*_1_(***X***, *U*) represents the baseline log-odds of the outcome as a function of covariates and the unmeasured confounder, and *g*_2_(***X***, *U*) represents the treatment effect on the log-odds scale. For settings with case1 = 0, the main effect is nonlinear; for settings with `case2 = 0`, the covariate-treatment interaction is nonlinear. Thus, by considering four settings, i.e.,(`case1, case2`): setting 1 (1, 1), setting 2 (1, 0), setting 3 (0, 1), and setting 4 (0, 0), we can test the performance of the proposed methods under both linear and nonlinear main effects/covariatetreatment interactions. In addition, in the generation of *T* and *Y*_*a*_, the unmeasured confounder *U* is associated with both, and thus the IV depending on *U* violates the core IV assumption and is indeed invalid. For all four setting, we fix *J* = 8, and vary the proportion of valid IVs in {100%, 90%}, the sample sizes *n* ∈ {400, 800, 1600}, and *ξ* ∈ {1.5, 3}, which leads to 48 scenarios in total.

We compare with multiple existing methods, including 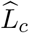 (with screening procedure) and 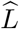 denoted as Baseline 1 and Baseline 2. Further, for the cases with possible invalid IVs, we generalize 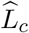 and 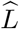, and combine them with our robust framework in Section 2.3, and we denote these methods as Baseline 1 and Baseline 2 as well. Besides, we also examine a naive approach (denoted as Naive method), where we take the exact maximum or minimum of the set of kernel estimates and construct a confidence interval only based on the variance of the maximizer or minimizer. The Naive method is expected to have lower coverage due to the post-selection bias. For each scenario, we first simulate samples as training data to fit all methods. For our proposed method and Baseline 2, the tuning parameters *γ* are selected from 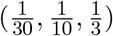 using five-fold cross-validation on the training dataset. To assess performance of different methods, we evaluate all methods on 25 testing samples (covariate values); for each sample, we compute the estimated lower and upper confidence bounds for the ITE and compare them with Baseline 1, Baseline 2, and Naive method. For all the methods, we implement multiplier bootstrap with 2000 repeats; for each simulation scenario, we repeat the entire procedure 200 times. We then calculate the empirical coverage probabilities w.r.t. *L* and *U*, or 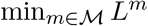 and 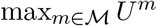 over the testing covariates and simulation replications.

Figure 1 summarizes the simulation results for two-sided 95% individual confidence bounds across the cases with and without invalid IVs in Settings (1,1) and (0,1); for invalid IV case, we apply our robust framework. The results for the remaining settings are reported in the online Supplementary Information. The top panel shows that our proposed method achieves the nominal coverage while the Naive method suffers from the post-selection bias, even when all IVs are valid; when there are invalid IVs, our proposed robust framework leads to valid inference procedures. The bottom panel shows that our method consistently identifies the largest fraction of individuals with significant treatment effects, which indicates that our proposed methods are indeed more efficient than the existing methods and their extensions, especially when there are invalid IVs. Overall, these results show that the proposed method achieves nominal coverage and yields higher efficiency even when invalid IVs exist.

**Figure 1:**
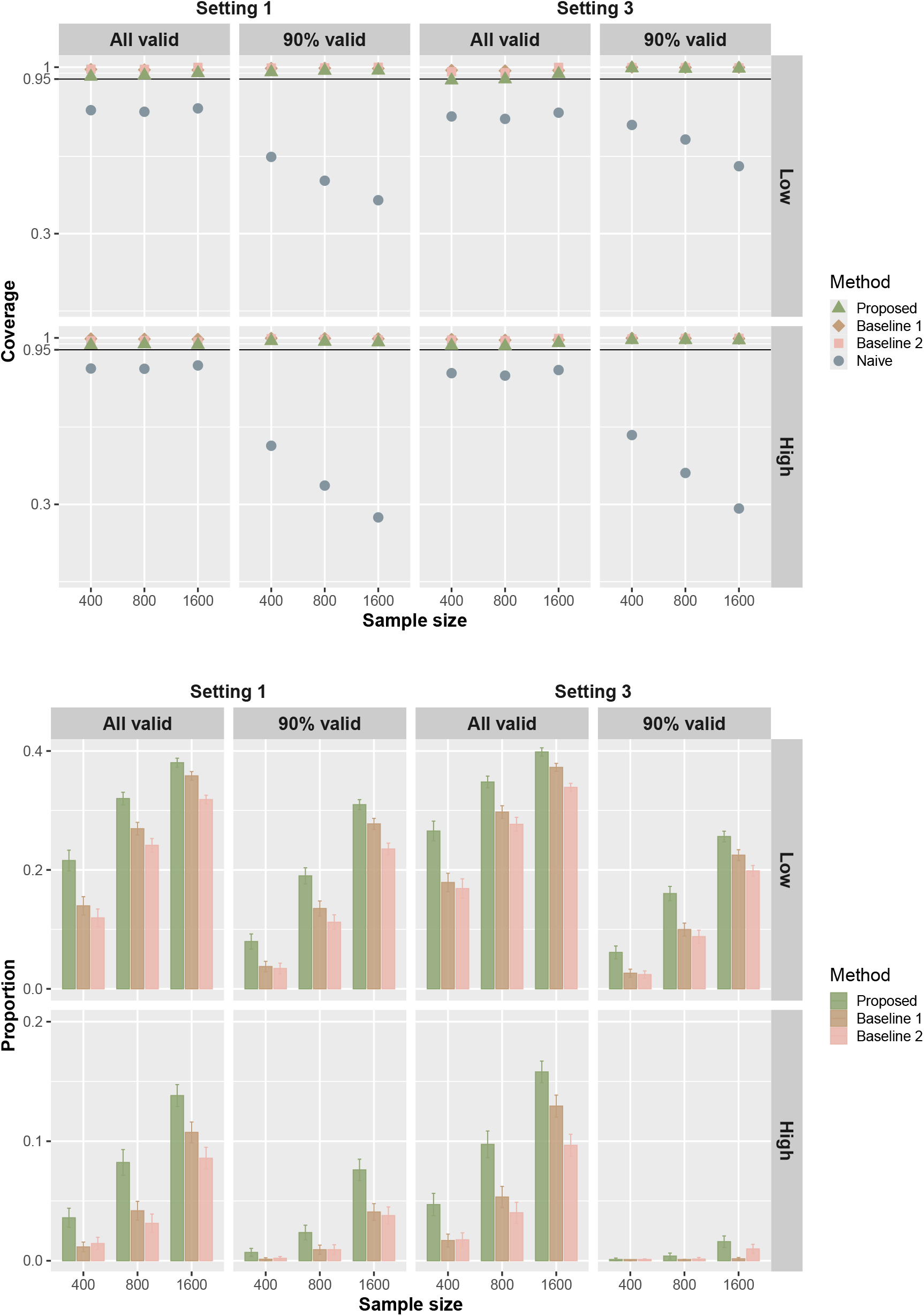
Empirical performance in Settings 1 and 3 over 200 repeats under low and high confounding effects (i.e., *ξ*). The top panel shows empirical coverage of nominal 95% confidence intervals, and the bottom panel shows the proportion of individuals whose confidence bounds exclude 0 (i.e., identified with nonzero causal effect).

## 5 Application to Estimating Genetic Effects in Alzheimer’s Disease

In this section, we apply our proposed methods to estimate heterogeneous genetic effects on AD development. We use the ADNI cohort, treating gene expression as the exposure and using genetic instruments to estimate individualized causal effects. The targeted gene in our case is TREM2, a highly potential AD risk gene [12]. Education levels are included as a covariate.

In our analysis, we use genotype data from the whole genome sequencing (WGS) for ADNI-1/GO/2 participants genotyped on the Illumina Omni2.5M array. The genotypes were processed with PLINK v2.0. The gene expression data were obtained from ADNI1/GO/2 Microarray whole blood samples. The phenotype data include the binary outcome of subjects’ baseline diagnosis as either AD or Control. To select IVs, we first implemented LD pruning by removing the variants that have pairwise *r*^2^ ≥ 0.01 to ensure independence between instruments, and kept those with minor allele frequency (MAF) ≥ 0.05. Then, we mapped cis-eQTLs with the R package MatrixEQTL using a ±1 Mb window around gene boundaries [32], and queried the GTEx portal to retrieve reported cis-eQTLs for specific genes. After mapping samples, we finally have a total sample size of *n* = 296, and we selected 6 cis-eQTLs as IVs. More details can be found in the online Supplementary Information.

In our analysis, we sample subjects from the entire dataset as testing samples. Then we conduct different methods on the entire dataset to construct confidence bounds for each individual in the testing samples. When implementing different methods, for our proposed methods and Baseline 2, the tuning parameters are selected among 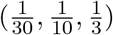. For all the methods, we implement multiplier bootstrap with 2000 repeats.

By applying our proposed methods and comparison methods, we have identified four subjects that have significant genetic effects when assuming all IVs are valid or 90% of IVs are valid. Table 1 shows an example for the subjects. The robust framework was applied to all methods when there are invalid IVs. When applying Baseline 1 and Baseline 2 to these subjects, the resulting confidence bounds are much wider, which shows that our proposed methods are more efficient than the comparison methods. In addition, these identified subjects are those with relatively low education levels, indicating that lower education might be associated with stronger evidence of a harmful effect of the target gene expression on AD risk. Figures 2 plots the averaged individual confidence bounds in the testing samples when education levels vary from low to high. It shows that individuals with low education have elevated risk; lower education levels might have a higher genetic risk of AD, i.e., the effect heterogeneity might be induced by education; and the pattern is consistent, even if assuming one IV is invalid.

**Table 1:** Individual confidence bounds with different methods and proportions of valid IVs.

| Subjects | Proposed |  | Baseline 1 |  | Baseline 2 |  |
| --- | --- | --- | --- | --- | --- | --- |
|  | all valid | 90% valid | all valid | 90% valid | all valid | 90% valid |
| Subject 1 | (0.941, 1) | (0.937, 1) | (0.938, 1) | (0.934, 1) | (0.455, 1) | (0.454, 1) |
| Subject 2 | (0.935, 1) | (0.928, 1) | (0.933, 1) | (0.925, 1) | (0.461, 1) | (0.458, 1) |

**Figure 2:**
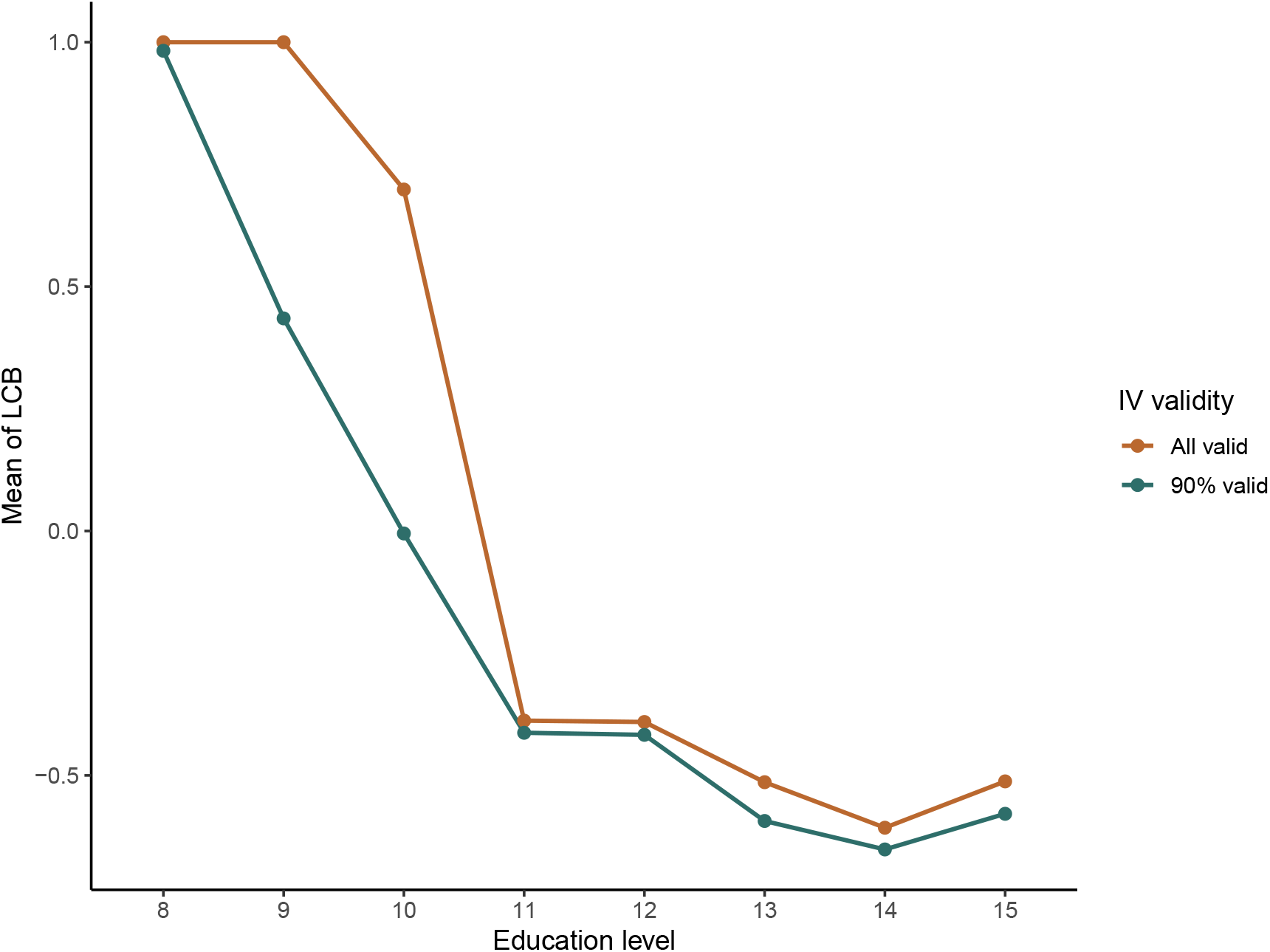
Mean lower confidence bound (LCB) of the individual causal risk using the proposed method across different educational levels, assuming all instruments are valid or allowing 10% invalid instruments.

## 6 Discussion

We develop a robust inference framework for ITE under MR that (i) requires minimal causal assumptions but is more efficient compared with existing literature and (ii) allows a controlled fraction of invalid IVs. Specifically, our approach combines nonparametric kernel regression with intersection-bound inference to deliver asymptotically sharp confidence intervals for individual-level causal bounds. Two adaptive steps are key: an estimator-specific variance standardization that leads to potentially tighter bounds and a bootstrap-based distributional shift that corrects selection bias and targets the true maximizers/minimizers. Building on this foundation, we construct an enumeration procedure that retains validity when some IVs may be invalid. Empirically, this strategy delivers more informative bounds relative to competing intersection-bound inference methods.

There are multiple extensions and future directions of our proposed methods. For example, focuses on the post-selection inference of subgroups identified in clinical trials. Our proposed method can be applied to this problem and provide more efficient inference for the treatment effects on the most effective subgroups. As another future direction, our current framework can be extended to high-dimensional settings with many candidate instruments and multiple exposures, such as multi-omics platforms. In such contexts, investigators often face multiple correlated biomarkers or gene expressions. Combining our approach with screening or representation learning tools may help manage this complexity while preserving individualized causal interpretations.

